# Opposite effects of lateralised transcranial alpha versus gamma stimulation on spatial attention

**DOI:** 10.1101/180836

**Authors:** Malte Wöstmann, Johannes Vosskuhl, Jonas Obleser, Christoph S. Herrmann

**Affiliations:** Department of Psychology, University of Lübeck, Lübeck, Germany; Experimental Psychology Lab, Center for Excellence “Hearing4all,” European Medical School, University of Oldenburg, Oldenburg, Germany

## Abstract

Spatial attention relatively increases the power of neural 10-Hz alpha oscillations in the hemisphere ipsilateral to attention. The functional roles of lateralised oscillations for attention are unclear. Here, 20 human participants performed a dichotic listening task under continuous transcranial alternating current stimulation (tACS) at alpha (10 Hz, vs sham) or gamma (47 Hz, vs sham) frequency, targeting left temporo-parietal cortex. Participants attended to four spoken numbers presented to one ear, while ignoring numbers on the other ear. As predicted, we found that alpha-tACS contralateral to the attended ear decreased recall of attended targets. Notably, gamma-tACS reversed the effect. Results provide a proof of concept that externally amplified oscillations can enhance spatial attention and facilitate attentional selection of speech. Furthermore, opposite effects of alpha versus gamma oscillations support the view that, across modalities, states of high alpha are incommensurate with active neural processing as reflected by states of high gamma.

## Introduction

When humans focus attention to one location in space, neural oscillatory alpha power (∼10 Hz) relatively increases in sensory areas in the hemisphere ipsilateral to the focus of attention and decreases in the contralateral hemisphere (1-3). The prevailing functional interpretation is that high ipsilateral alpha power inhibits cortical activity in the hemisphere processing the unattended side of space, which agrees with the functional inhibition theory of alpha oscillations (4, 5). This view receives further support by studies showing that gamma power (>40 Hz), which reflects active cortical processing (6) and correlates negatively with alpha power (7), lateralises in the opposite way compared to alpha power during spatial attention (8-10). In the present combined behavioural and tACS (transcranial alternating current stimulation) study, we test whether lateralised alpha and gamma oscillations are just epiphenomena of attention or whether they, when externally amplified, modulate accuracy of auditory spatial attention to speech.

In the visual modality, rhythmic transcranial magnetic stimulation (rTMS) at alpha frequency applied to one hemisphere has been shown to enhance perception (11) and to improve memory for targets ipsilateral to the stimulated hemisphere (12, for a review of TMS effects on visuospatial attention, see 13). In audition, however, the existence of an *auditory alpha rhythm* (14) and functional roles thereof are notoriously more challenging to assess experimentally (15, 16). Also, the link between the amplitude of lateralised auditory alpha oscillations and spatial attention is purely correlational so far: In a recent magnetoecephalograhphy (MEG) study (17), we found that stronger temporally modulated alpha lateralisation (i.e., high ipsi-and low contralateral alpha power) predicted better recall of attended speech in a dichotic listening task. Here, we use the same task and aim to boost participants’ endogenous neural oscillations using tACS to increase the amplitude of alpha or gamma oscillations (see e.g., 18, 19, 20) in a superior temporal/inferior parietal cortex region in the left hemisphere. This region was found to exhibit marked and directly performance-related alpha lateralisation during this task (17).

If lateralised oscillatory power were a mere *correlate* or epiphenomenon of neural processes that instantiate attention, alpha- and gamma-tACS should not affect accuracy of spatial attention. However, our findings demonstrate that left hemispheric alpha-versus gamma-tACS do modulate oppositely the recall of attended speech on the left versus right side. This is evidence that lateralised oscillations are a functionally significant *substrate* of spatial attention.

## Results & Discussion

In a dichotic listening task, *n* = 20 participants attended to four spoken numbers presented to one ear, while they ignored four numbers presented simultaneously to the other ear. In the end of each trial, participants had the task to select the four numbers presented to the to-be-attended side from a visually presented number pad (Fig. 1A). This paradigm has been shown to induce robust lateralisation of neural oscillations, with relative high alpha power in auditory and parietal cortex regions in the hemisphere ipsi-versus contralateral to the focus of attention (Fig. 1B, left; 17). Here, we aimed at increasing the amplitude of participants’ lateralised oscillations during dichotic listening using left-hemispheric tACS at alpha (10 Hz) or gamma (47.1 Hz) frequency.

**Figure 1.**
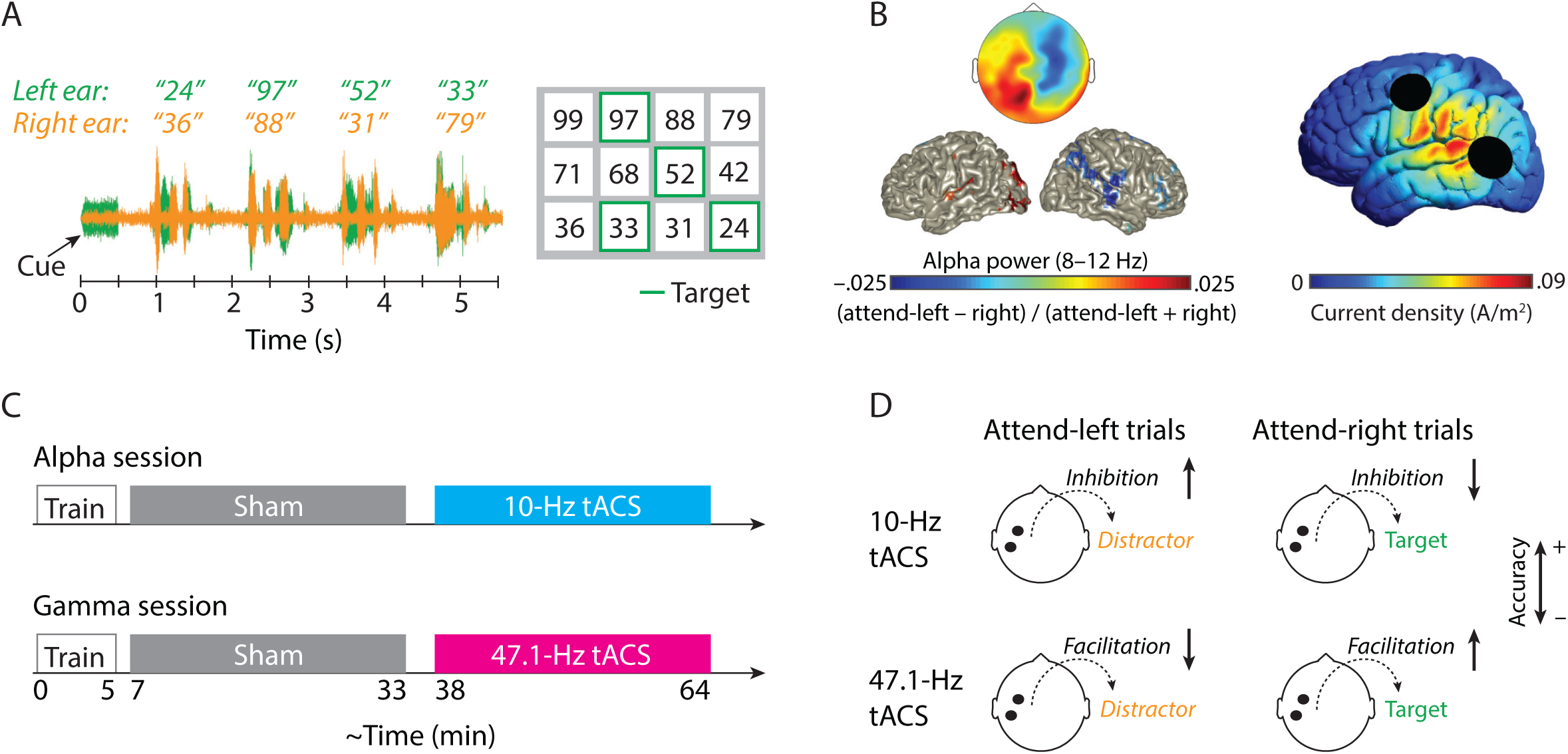
(A) Dichotic listening task. A cue tone on one ear (left in this example) indicated the to-be-attended side on this trial. Four spoken numbers were presented to the left ear, and four different (same-talker) numbers to the right ear. The task was to select the four to-be-attended numbers (i.e., targets) from a visually presented number pad shown after the end of auditory stimulation. Green edges, which were not shown to participants, indicate correct responses on this trial. (B, left) Topography and brain surfaces show hemispheric lateralisation of alpha power (8–12 Hz) during the presentation of numbers, obtained in a previous magnetoencephalography (MEG) study using the same task (17) by contrasting attend-left versus attend-right trials: (attend-left–attend-right) / (attend-left+attend-right). Overlays on brain surfaces are masked at *p* = 0.05; uncorrected. (B, right) Simulation of current densities emitted by our tACS stimulation setup as calculated in SimNIBS (21). Stimulation was intended to target temporo-parietal regions (around traget region pSTG) in the left hemisphere. (C) The experiment was divided in two sessions, which took part on different days. In each session, participants first performed a short training, followed by a sham run of the experiment without tACS stimulation, a short break, and finally another run of the experiment under continuous tACS stimulation lasting 25 min (alpha session: 10 Hz; gamma session: 47.1 Hz). (D) tACS stimulation was applied to electrodes TP7 and FC5 (black dots on topographies and in B, right). Alpha-versus gamma-tACS were expected to induce differential patterns of increasing (upward pointing arrows) and decreasing accuracy (downward pointing arrows) in attend-left/-right trials (see text for details).

In humans, left auditory cortex regions receive input predominantly from the right ear and vice versa for right auditory cortex (22, 23). Therefore, our tACS setup, which targeted primarily left auditory and parietal regions (Fig. 1B, right), was expected to affect neural processing of the right-ear input in a highly predictable way (Fig. 1D): Left-hemispheric alpha-tACS should inhibit neural processing of the right-ear input, leading to reduced accuracy in attend-right but enhanced accuracy in attend-left trials. Contrary, left-hemispheric gamma-tACS should facilitate neural processing of the right-ear input, leading to enhanced accuracy in attend-right but reduced accuracy in attend-left trials.

### Right-ear advantage in the dichotic listening task

For statistical analyses, we used linear mixed-effects models to regress the proportion of correctly recalled digits in each trial on the predictor variables session (alpha vs gamma), stimulation (sham vs tACS), and to-be-attended side (left vs right). As expected for dichotic listening, proportion correct was higher for attend-right than attend-left trials (*b* = 0.075; *F*_*1*, *8773*_ = 115.15; *p* < 0.001). This demonstrates the well-known right-ear advantage (REA; 24), which rests on structural (25), but also attentional hemispheric asymmetries (26).

Since the right-ear advantage in dichotic listening tasks is known to be of limited test–retest reliability in the individual (27, 28), we included a sham run immediately before the tACS run in each experimental session (see Fig. 1C). Thus, tACS effects could be quantified by contrasting each participants’ tACS run with the sham run of the respective session.

### Alpha- and gamma-tACS differentially modulate auditory spatial attention

Most importantly, and directly in line with our hypotheses (Fig. 1D), the session × stimulation × to-be-attended side interaction was significant (*b* = 0.062; *F*_*1, 8773*_ = 11.68; *p* < 0.001). Compared to sham, left-hemispheric alpha-tACS enhanced the proportion correct in attend-left trials but decreased the proportion correct in attend-right trials (Fig. 2A; stimulation × to-be-attended side interaction: *b* = –0.024; *F*_*1, 4377*_= 3.57; *p* = 0.059), while the pattern of results precisely reversed for gamma-tACS (Fig. 2B; stimulation × to-be-attended side interaction: *b* = 0.038; *F*_*1, 4377*_ = 8.83; *p* = 0.003).

**Figure 2.**
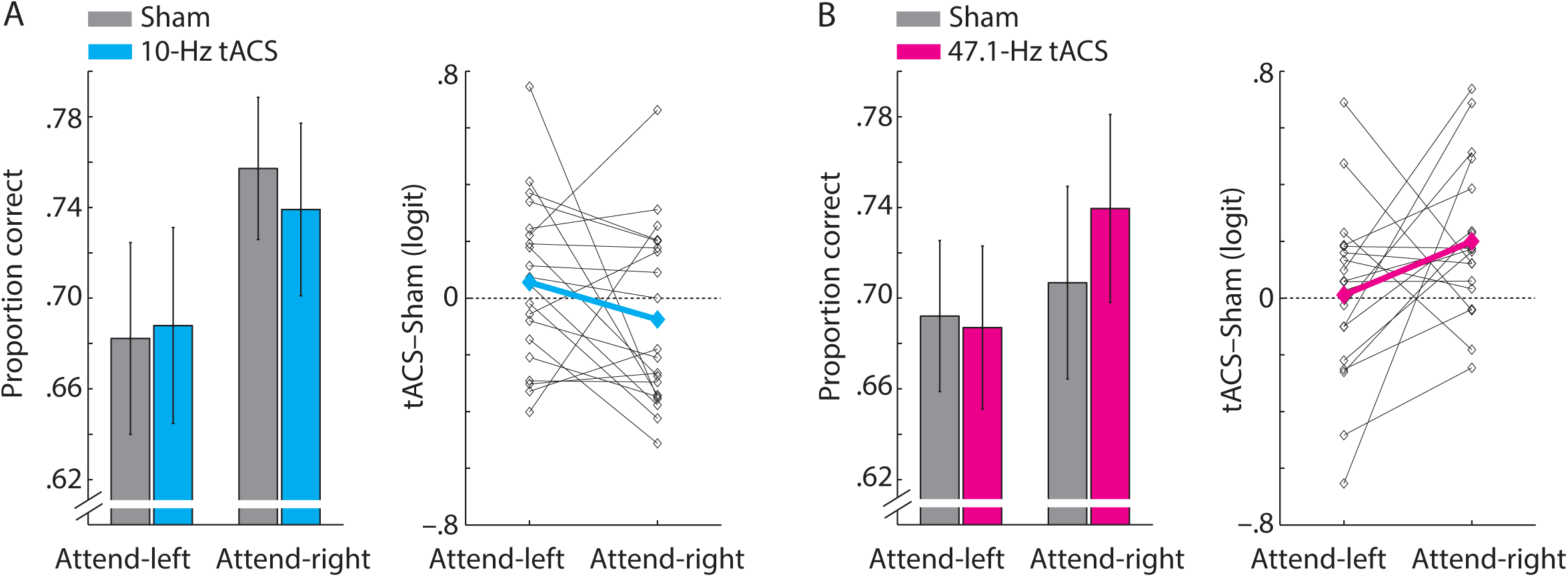
Effects of left-hemispheric tACS on behaviour in the dichotic listening task. (A) Results of the alpha session (10-Hz stimulation). Bars and errorbars show mean ±1 between-subject SEM of proportion correct responses in the dichotic listening task for attend-left and attend-right trials. The line plot shows the tACS effect (10-Hz tACS–Sham; logit-transformed) for *n* = 20 individual participants (thin lines), bold line shows average across participants. (B) Same as A but for the gamma session (47.1-Hz tACS).

So far, electric stimulation studies targeting one hemisphere have found shifts of visuo-spatial attention in case of transcranial direct current stimulation (tDCS; 29, 30), whereas tACS at alpha frequency has produced inconclusive (31) or non-replicable results (32). For auditory spatial processing, one study aimed at modulating the right-ear advantage using tDCS applied to either the left or right hemisphere during dichotic listening but did not observe hemisphere-specific performance modulations (33). To the contrary, the opposing effects of left-hemispheric alpha-versus gamma-tACS observed here show that externally amplified lateralised oscillations modulate the focus of auditory attention in a frequency-specific way: tACS-increased alpha oscillations inhibit, while tACS-increased gamma oscillations facilitate, the attentional selection of speech presented contralateral to the stimulated hemisphere. Thus, both lateralised alpha and gamma oscillations exert a functional relevance in auditory spatial attention.

Our participants were stimulated continuously for 25 minutes and, since the trial timing was not fixed but depended on a participant’s response speed, the onsets of auditory events (cue tone and numbers) were randomly distributed across the cycle of the stimulated alpha or gamma oscillation. Thus, whereas previous research found phase effects of alpha-tACS on auditory perception (e.g., 34), the present study demonstrates that tACS modulates spatial attention to auditory events which are non-phase-locked to the stimulated oscillation.

Based on the theoretically proposed (e.g., 4) and empirically observed opposing roles of inhibitory alpha and facilitatory gamma power for spatial attention (e.g., 35), we expected hemispheric gamma-tACS to reverse the effect of alpha-tACS. Our results confirm this. Previous tACS studies have shown that high-frequency random noise stimulation (100–640 Hz) targeting auditory cortex enhanced auditory responses in the EEG (36) and improved auditory gap-detection performance (37). Together with these studies, our results support the view that gamma-tACS can facilitate auditory processing and thus affects auditory spatial attention inversely compared to tACS-entrained inhibitory alpha oscillations.

### Left-hemispheric tACS affects attend-right trials

Separate analyses of attend-left and attend-right trials revealed that the session (alpha vs gamma) × stimulation (tACS vs sham) interaction was significant for attend-right trials (*b* = 0.051; *F*_*1, 4377*_ = 17.41; *p* < 0.001) but not for attend-left trials (*b* = –0.011; *F*_*1, 4377*_ = 0.67; *p* = 0.413). We assume that this is due to our left-hemispheric locus of stimulation, which, due to the contralateral organization of the human auditory system, modulated auditory processing of task-relevant target speech in attend-right trials but processing of task-irrelevant distractor speech in attend-left trials (see Fig. 1D). Recall accuracy in the dichotic listening task likely depends more on attending to target speech, and hinges only indirectly on ignoring the distractor (for a similar argument, see 38). Thus, the impact of tACS-increased oscillations of either frequency on target speech (in attend-right trials) was larger than the impact of tACS-increased oscillations on distractor speech (in attend-left trials).

Note that stronger effects of left-hemispheric alpha- and gamma-tACS in attend-right trials indirectly support the feasibility of our stimulation setup in targeting primarily regions in the left hemisphere. In theory, right-instead of left-hemispheric tACS in our dichotic listening task should induce stronger performance modulations in attend-left trials. But functional asymmetries of left versus right auditory cortex regions could further complicate this reasoning (39, 40).

### Functional roles of lateralised alpha and gamma oscillations

The pattern of our results is in striking accordance with the hypothesised roles of lateralised inhibitory alpha and facilitatory gamma oscillations for spatial attention (Fig. 1D). It is important to note, however, that even experiments employing perturbation techniques such as tACS allow only limited causal inference (41). For instance, it might be that tACS-increased lateralised alpha oscillations causally inhibit neural processing in the stimulated brain regions, whereas tACS-increased gamma oscillations are causally ineffective by themselves but instead provide a means to decrease the power of causally effective alpha oscillations (for an effect of gamma-tACS on alpha oscillations, see 42).

Furthermore, our results do not reveal at which level of auditory processing tACS-increased alpha and gamma oscillations are effective. In line with the predominant views on the functional roles of neural oscillations (e.g., 43), it is reasonable to assume that alpha-tACS affects behaviour through inhibition of (top-down) attention to one side of space whereas gamma-tACS affects behaviour through facilitation of neural (bottom-up) encoding (see e.g., 10) of auditory stimuli in left versus right auditory cortex. Alternatively, alpha- and gamma-tACS might affect the same neural processes but in opposite direction, but this view is somewhat weakened by the present data: the effects of alpha- and gamma-tACS on the recall of target numbers (i.e., tACS–sham, logit transformed) were not significantly negatively correlated (attend-left trials: *r* = 0.19; *p* = 0.427; attend-right trials: *r* = 0.02; *p* = 0.938). Combining brain stimulation, neuroimaging and behavioural measures should test these alternative explanations in future studies (44).

### Conclusions

Lateralised neural oscillations across the cerebral hemispheres are a well-known signature of spatial attention. Here, we show that increased lateralised alpha and gamma oscillations using transcranial alternating current stimulation hold the power to modulate spatial attention to one of the most important sensory signals in human environments, that is, speech. In agreement with prevailing views on alpha and gamma oscillations, our results support the functional relevance of these regimes of neural oscillations for auditory spatial attention: The attentional selection of speech presented to the left or right side is inhibited when alpha power increases in the contralateral hemisphere (and vice versa for gamma power).

## Materials and Methods

### Participants

Twenty young healthy participants (19–31 years, 10 females) with no history of neurological or psychiatric disorders participated. One participant was not fully right handed (laterality quotient of 20 on a scale from –100 (left handed) to +100 (right handed)) according to the Edinburgh inventory (45). Two participants were non-German native speakers but of sufficient German language proficiency to perform the dichotic listening task. Experimental procedures were approved by the local ethics committee of the University of Oldenburg (Komission für Forschungsfolgenabschätzung und Ethik).

### Auditory stimuli

We used German, female-voiced 4-syllable numbers from previous studies (46, 47). Recordings of spoken numbers from 21 to 99 (excluding integer multiples of 10) with a sampling rate of 44.1 kHz were shortened in Praat (version 6.0.14) by a factor of 0.85, resulting in a mean (±SD) number duration of 0.96 s (±0.05 s). We determined the perceptual center (P-center) of each number (48) as the time point when the number signal’s broad-band envelope (15-Hz lowpass-filtered modulus of the Hilbert transform) reached 50% of the first syllable’s peak amplitude. In the following, the onset of a number refers to its perceptual center. For the spatial cue, we used a monaural 1000-Hz sine tone of 500-ms duration.

### Dichotic listening task

The dichotic listening task was a slightly speeded version of a task used previously (17). For each trial, eight different numbers were selected randomly from the pool of all numbers; four to-be-presented to the left, the other four to-be-presented simultaneously to the right ear. For simultaneously presented numbers on the left and right ear, perceptual centers were temporally aligned and numbers were distinct in their first and second digit (e.g., co-occurences of “35” and “37” or “81” and “21” were avoided).

Each trial started with the presentation of the cue (to one ear) to indicate whether participants had to attend to the left or right in this trial. The cue was followed (after 500 ms) simultaneously by four spoken numbers presented to the left ear and four different numbers to the right ear (Fig. 1A). The onset-to-onset time interval of two subsequent numbers was 1.25 s. Presentation of acoustic stimuli (cue and numbers) took on average (±SD) 5.66 s (± 0.04). Finally, cue tone and spoken numbers were embedded in continuous background noise (white noise; +10 dB SNR). Cue tone and background noise had 50-ms linear onset and offset ramps.

Approximately 0.5 s (jittered 0.3 – 0.7 s) after the offset of the last two simultaneous numbers a response screen was shown that contained 12 numbers (four from the to-be-attended side, four from the to-be-ignored side, and four random numbers not presented on any side). Participants were asked to use a mouse to select the four numbers that appeared on the to-be-attended side in any order. Numbers on the response screen were presented in either ascending or descending order (randomised from trial to trial) to prevent motor preparation during a trial. After selection of four numbers, the next trial started automatically (after approximately 1 s, randomly jittered 0.8 – 1.2 s). Auditory materials were presented via Sennheiser HD 25-1 II headphones. In one run of the experiment a participant performed 110 trials, which took on average 26’31’’ (±2’4’’ SD) to complete. Trial order was fully randomised with the constraint that the spatial cue appeared on the left side in half of the trials in each run.

### tACS stimulation

We used a lateralised tACS stimulation setup, which was designed to target left-hemispheric posterior superior temporal gyrus (pSTG) and surrounding auditory and parietal cortex regions. Round electrodes of 3 cm diameter were placed at sites FC5 and TP7 according to the international 10-10 system. Electrode impedance was kept below 10 kΩ. The stimulator (DC stimulator plus, Eldith, 419 NeuroConn, Ilmenau, Germany) emitted a sinusoidal alternating current with no DC offset at frequencies of either 10 Hz (alpha) or 47.1 Hz (gamma) at a strength of 1 mA (milliampere) peak-to-peak. The gamma stimulation frequency of 47.1 Hz was chosen to not match a multiple (i.e., harmonic) of the alpha frequency. Stimulation amplitude was ramped up and down linearly over 20 or 94.2 cycles (2 Sec) for 10 Hz and 47.1 Hz stimulation frequencies, respectively. Overall tACS was applied continuously for 25 minutes between the two ramping periods.

For sham stimulation, the tACS setup was the same as for alpha and gamma stimulation, except that the stimulation consisted only of the ramping periods at the beginning and subsequently remained off for the rest of the run.

### Procedure

In total, each participant performed four runs of the dichotic listening task (resulting in 4 × 110 = 440 trials per participant), separated in two sessions taking place on different days: alpha session (sham run & alpha-tACS run) and gamma session (sham run & gamma-tACS run). Session order (alpha-->gamma vs gamma--> alpha) was balanced across participants and the two sessions were separated by 5–14 days (average: 7.58 days). Each session started with a short training (approx. 5 min) to familiarise the participant with the task procedure. Next, participants performed the sham run, followed by a short break and the alpha/gamma-tACS stimulation run. Within each session the sham run was tested first in order to avoid possible confounds of tACS after-effects, which are known to outlast the duration of stimulation considerably (49, 50).

### Statistical analyses

The major dependent measure in the present study was the proportion of correctly recalled numbers on attend-left versus attend-right trials. On each trial participants selected four numbers from the response screen, resulting in five possible proportions of correctly recalled numbers: 1 (four out of four), 0.75 (three out of four), 0.5 (two out of four), 0.25 (one out of four), 0 (zero out of four).

The study implemented a 2 (session: alpha vs gamma) × 2 (stimulation: sham vs tACS) × 2 (to-be-attended side: left vs right) within-subject design. For the statistical analysis we fitted linear mixed-effects models using the *lme4* package for R (version 2017-03-06) and Rstudio (version 1.0.136). In essence, participants’ single-trial data were used to model the response variable proportion correct on the three predictors. To obtain *p*-values for predictors (51), we used the *anova* function using Satterthwaite approximation implemented in the *lmerTest* package. These *p*-values were also compared to *p*-values obtained with the *mixed* function using parametric bootstrapping implemented in the *afex* package, which yielded almost identical results.

For visualization of group data (bar graphs in Fig. 2), we averaged the proportion correct for each experimental condition. For visualization of single-subject data (line graphs in Fig. 2), we logit-transformed the proportion correct data for each subject and condition (52), followed by calculation of the tACS effect (tACS–sham), separately for the alpha and gamma session.

### Side effects of tACS

According to questionnaires answered in the end of each session (alpha & gamma), the side effects reported by most of our 20 participants were tingling (alpha: 18; gamma: 15), difficulty in concentration (alpha: 17; gamma: 15), and tiredness (alpha: 12; gamma: 10). However, only a subset of participants attributed these side effects to the tACS stimulation (tingling, alpha: 13, gamma: 12; difficulty in concentration, alpha: 7, gamma: 6; tiredness, alpha: 5, gamma: 4). Intensities of these side effects (rated on a scale from 0 = ‘no’ to 4 = ‘strong’) did not differ between alpha and gamma sessions (Wilcoxon signed rank tests; all *p* > 0.1).

Participants were also asked in the end of each session to indicate for individual runs of a session (sham & tACS) whether they think they were stimulated. For the alpha session, 7 participants repoted stimulation during sham and 12 during tACS (McNemar test; *p* = 0.18). For the gamma session, 6 participants reported stimulation during sham and 15 during tACS (*p* = 0.012). Stronger sensation of stimulation in the gamma session might explain a change in overall performance but not our specific hypothesised response patterns (i.e., differential performance modulation in attend-left versus attend-right trials; see Fig. 1D). We thus consider these side effects uncritical to the results of the present study.

### Statistical control analyses

The major result of this study was the significant session × stimulation × to-be-attended side interaction, obtained via mixed-effects linear regression analysis. We used two additional linear mixed-effects models to confirm the robustness of this interaction. First, this interaction remained significant when we controlled for session order (alpha-->gamma vs gamma-->alpha; *b* = 0.062; *F*_*1, 8766*_= 11.71; *p* < 0.001) and laterality quotient (*b* = 0.062; *F*_*1, 8766*_= 11.72; *p* < 0.001), assessed by the Edinburgh handedness questionnaire (45).

Second, since non-normality of residuals is a potential confound in linear regression, we backed-up our statistical analysis using Poisson regression to model the number of errors on each trial (0, 1, 2, 3, or 4), which confirmed the significant right-ear advantage (*b* = –0.003; *F*_*1*_ = 70.07; *p* < 0.001) and session × stimulation × to-be-attended side interaction (*b* = –0.002; *F*_*1*_ = 7.94; *p* = 0.005). Third, in addition to the single-trial mixed effects approach used in the present study, we also performed a traditional repeated-measures ANOVA on the average proportion of correctly recalled numbers in each condition, which also revealed a significant right-ear advantage (*F*_1, 19_= 13.23; *p* = 0.002; *η*^*2*^_*P*_ = 0.41) and session × stimulation × to-be-attended side interaction (*F*_1, 19_ = 7.63; *p* = 0.012; *η*^*2*^_*P*_ = 0.29).

Furthermore, it is critical in the present study to control for possible changes in proportion correct across the time of an experimental run, for three reasons: First, our ethics approval allowed 25 min of continuous tACS stimulation, but some participants required >25 min to complete a run of the experiment (mean run duration ±1SD; alpha session, sham: 27’14’’ ±2’53’’, tACS: 26’25’’ ±3’11’’; gamma session, sham: 26’48’’ ±2’59’’, tACS: 25’37’’ ±3’05’’). Second, in each session, we first measured the sham run followed by the tACS stimulation run, leading to significantly faster completion of tACS compared to sham runs (repeated-measures ANOVA; *F*_1,19_= 13.81; *p* = 0.001; *η*^*2*^*P* = 0.42). Third, it has been speculated that during longer time periods of continuous lateralised tACS stimulation, the applied current would initially reach target areas and later spread to non-target areas in the contralateral hemisphere, which could confound effects of lateralised tACS (31). For these reasons, we conducted two additional linear regressions to control for single-trial offset times (measured in milliseconds relative to the offset of the ramp-up of the stimulation amplitude in the beginning of each run) and stimulation offset, coded 0 for all trials completed within the first 25 min and 1 for all trials completed thereafter. Importantly, the session × stimulation × to-be-attended side interaction was significant when we controlled for single-trial offset time (*b* = 0.063; *F*_*1*, 8766_= 12.12; *p* < 0.001) as well as stimulation offset (*b* = 0.06; *F*_*1*, 8766_= 5.64; *p* = 0.018). Furthermore, single-trial offset time and stimulation offset did not exhibit main or interaction effects on proportion correct (all *p* > 0.05).

### Effect sizes

Since there are no standard effect size measures for individual predictors and their interactions in linear mixed models, we report the unstandardized coefficient (*b*), which corresponds to the estimated change in the dependent variable (proportion correctly recalled numbers) when the predictor increases by one level. For repeated-measures ANOVAs, we report partial eta squared (*η*^*2*^_*P*_).

## Acknowledgements

We thank Erika Puiutta and Tania Pollok for the data acquisition and Sarah Tune for her help with the statistical analysis.

